# VEHiCLE: a Variationally Encoded Hi-C Loss Enhancement algorithm

**DOI:** 10.1101/2020.12.07.413559

**Authors:** Max Highsmith, Jianlin Cheng

## Abstract

Chromatin conformation plays an important role in a variety of genomic processes. Hi-C is one of the most popular assays for inspecting chromatin conformation. However, the utility of Hi-C contact maps is bottlenecked by resolution. Here we present VEHiCLE, a deep learning algorithm for resolution enhancement of Hi-C contact data. VEHiCLE utilises a variational autoencoder and adversarial training strategy to enhance contact maps, making them more viable for downstream analysis. VEHiCLE expands previous efforts at Hi-C super resolution by providing novel insight into the biologically meaningful and human interpretable feature extraction. Using a variational autoencoder VEHiCLE provides a user tunable, full generative model for generating synthetic Hi-C data while also providing state-of-the-art results in enhancement of Hi-C data across multiple metrics.

## Introduction

Hi-C data, an extension of chromosome conformation capture assay (3C) is a biological assay which can be used to inspect the three-dimensional (3D) architecture of a genome ^1^. Hi-C data can be used for downstream analysis of structural features of chromosomes such as AB compartment, Topological Associated Domains (TADs), loops, and 3D models. Changes in chromosomal conformation have been empirically demonstrated to impact a variety of genomic processes including gene methylation and gene expression ^2^.

When analysing Hi-C data, reads are usually converted into contact matrices, where each cell entry corresponds to the quantity of contacts between the two regions indexed by row and column ^3,4^. The size of an individual region in this contact matrix is referred to as the resolution or bin size^4^. The resolution of a contact matrix is usually selected based on the quantity of read pairs in an individual Hi-C experiment, with a higher quantity of read pairs permitting a higher resolution. Certain genomic features, such as TADs, can only be meaningfully identified using high resolution contact matrices, however if a matrix resolution is selected with insufficient read coverage the matrices can be overly sparse. One method to address this issue is to run additional Hi-C experiments, however because of experimental costs this is not always a feasible solution.

To solve this problem previous groups have utilized methods from the field of Image super-resolution to improve Hi-C contact matrix resolution. The first of these networks was HiCPlus ^5^, a simple neural network optimized using mean squared error (mse). HiCPlus was then improved upon by HiCNN ^6^ by adjusting network architecture. Next hicGAN ^7^ was proposed, introducing the use of Generative Adversarial Networks (GAN), which generated high resolution contact maps conditioned on low resolution input. The network DeepHiC ^8^ maintained the GAN loss function while extending it to also include a perceptual loss function derived from VGG-16 trained on image data. The model HiCSR ^9^ continued the advancement by introducing the use of a deep autoencoder as a feature extraction mechanism.

Our network, the **V**ariationally **E**ncoded **Hi**-**C L**oss **E**nhancer (VEHiCLE), extends the approach of conditional generative adversarial networks by using an integrated training approach inspired by literature in the domains of deep learning and genomics. First, VEHiCLE incorporates a variational autoencoder which extracts biologically meaningful features from Hi-C data. Second, VEHiCLE’s decoder network is engineered to provide an easy to use generative model for Hi-C data generation which smoothly maps user tunable, low dimensional vectors to Hi-C contact maps independent of any low sampled input. Third, VEHiCLE incorporates a biologically explicit loss function based on Topologically Associated Domain identification to ensure accurate downstream genomic analysis.

VEHiCLE obtains state of the art results in the task of Hi-C superresolution across a variety of metrics pulled from the domains of Image analysis and Hi-C quality/reproducibility. VEHiCLE enhanced data show successful retrieval of important downstream structures such as TAD identification and 3DModel generation while also providing novel human interpretability of its enhancement process.

## Approach

### Description of VEHiCLE network training

Vehicle is trained as an adversarial network conditioned on low resolution input. The network is trained using a composite loss function made up of 4 sub loss functions: Adversarial loss, Variational loss, MSE loss, and Insulation loss. An overview of the training mechanism is displayed in figure 1a. The intellectual motivation for each of these loss functions is outlined below.

**Figure 1.**
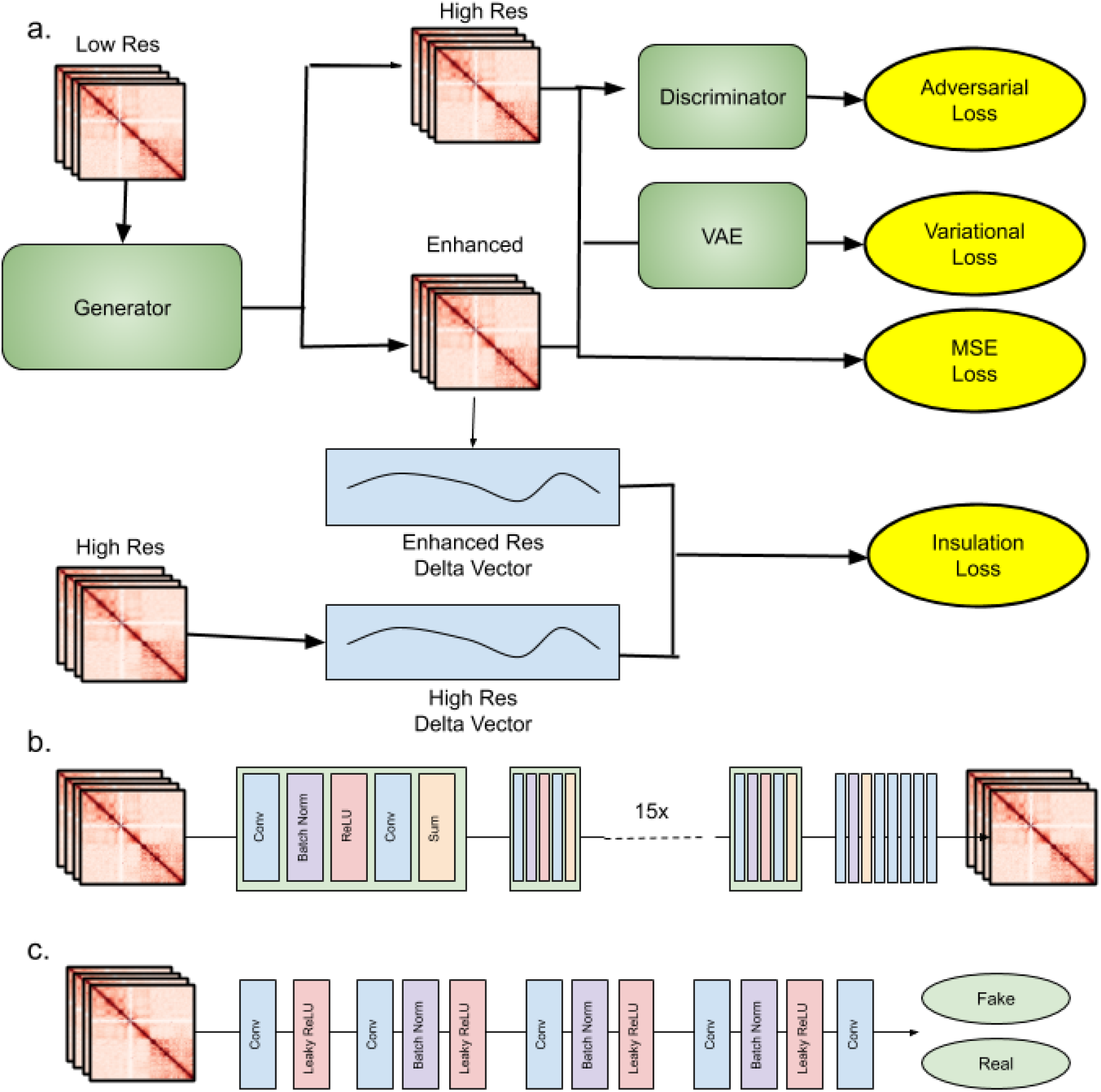
VEHiCLE Architecture (a) overview of training strategy (b) generator architecture (c) Discriminator architecture.

#### Adversarial loss function

Generative adversarial networks (GANs) are a popular deep learning based framework for generative modeling which has gained traction in a wide variety of tasks including image superresolution. GANs were first introduced to the field of Hi-C super resolution through hicGAN, and later improved upon in DeepHiC and HiCSR. A GAN uses two key networks: a generator *G* (figure 1b) and a discriminator *D* (figure 1c). The generator takes samples from an input distribution and generates enhanced matrices. The Discriminator is trained on a collection of inputs including real high resolution Hi-C samples as well as enhanced resolution Hi-C samples, and attempts to determine whether individual samples are real or enhanced. The two networks are trained in a game where the generator is rewarded for successfully tricking the discriminator and the discriminator tries to minimize classification mistakes.

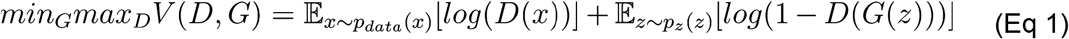

The generator loss function is defined as:

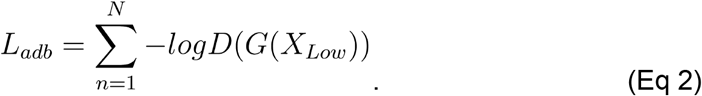

#### Variational Loss

Autoencoders are deep learning systems which map inputs from sample space to a condensed latent space via an encoder and then reconstruct images in sample space from the latent space using a decoder. The use of autoencoders for the task of Hi-C data super resolution was originally proposed in our preprint ^10^ for the task of denoising Hi-C data. They were then suggested by Dimmick et al ^9^ as tools for training super resolution networks by using the features extracted by passing Hi-C data through a trained autoencoder as a loss function. In this manuscript we expand upon this strategy, but replace their network with a different flavor of network called the variational autoencoder ^11^.

Similar to vanilla autoencoders, variational autoencoders (VAE) aim to condense data into lower dimensional space, however they have the advantage of providing smooth feature representation which can permit the construction of powerful generative models. To obtain these advantages VAE relly upon a statistical method called variational inference^11^. This method frames the tasks of encoding and decoding as an ancestral sampling problem with two steps: First, a latent variable *z* is sampled from a prior distribution *P_θ_*(*z*). Second, the observed variables *x* are drawn from a likelihood distribution *P_θ_*(*x*|*z*).

To encode the observed variable *x* requires the computation of the posterior distribution *P_θ_*(*z*|*x*). However because this is computationally intractable, instead one approximates the posterior by choosing a parametric family of recognition models *q*_Φ_(*z*|*x*) and selects parameters that minimize the divergence between our recognition model and the true underlying distribution via a probabilistic dissimilarity metric called KL-divergence,

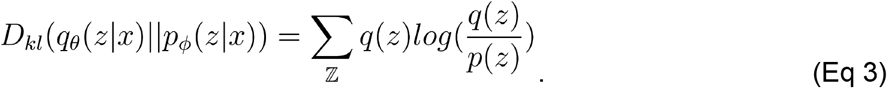

By performing some algebra outlined in Kingma and welling ^11^ variational autoencoders are trained using the following loss function

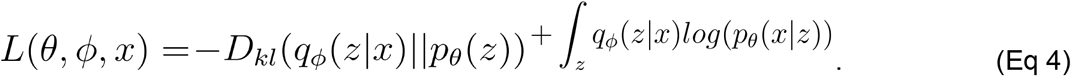

The integral term on the far right of the loss function ensures that the reconstruction outputs of our networks are highly similar to their original inputs, while the KL divergence term causes the latent space distribution of values to closely resemble a vector of gaussian random variables . This imposition of gaussian similarity on the latent space results in advantages in quality of extracted features and the procurement of a generative model.

To create the variational loss function we first train our variational autoencoder using high resolution contact matrices as both inputs and labels. In each experiment our VAE network is trained using the same chromosomes as the overall VEHiCLE network. The variational autoencoder maps vectors from data space into condensed latent space, which we interpret as a lower dimensional feature vector (Fig2 ab). Because the variational autoencoder training strategy imposes a gaussian distribution of the latent space variables and because our decoder maps latent vectors back into data space in a relatively smooth manner we expect highly similar Hi-C contact matrices to contain similar latent space profiles.

**Figure 2.**
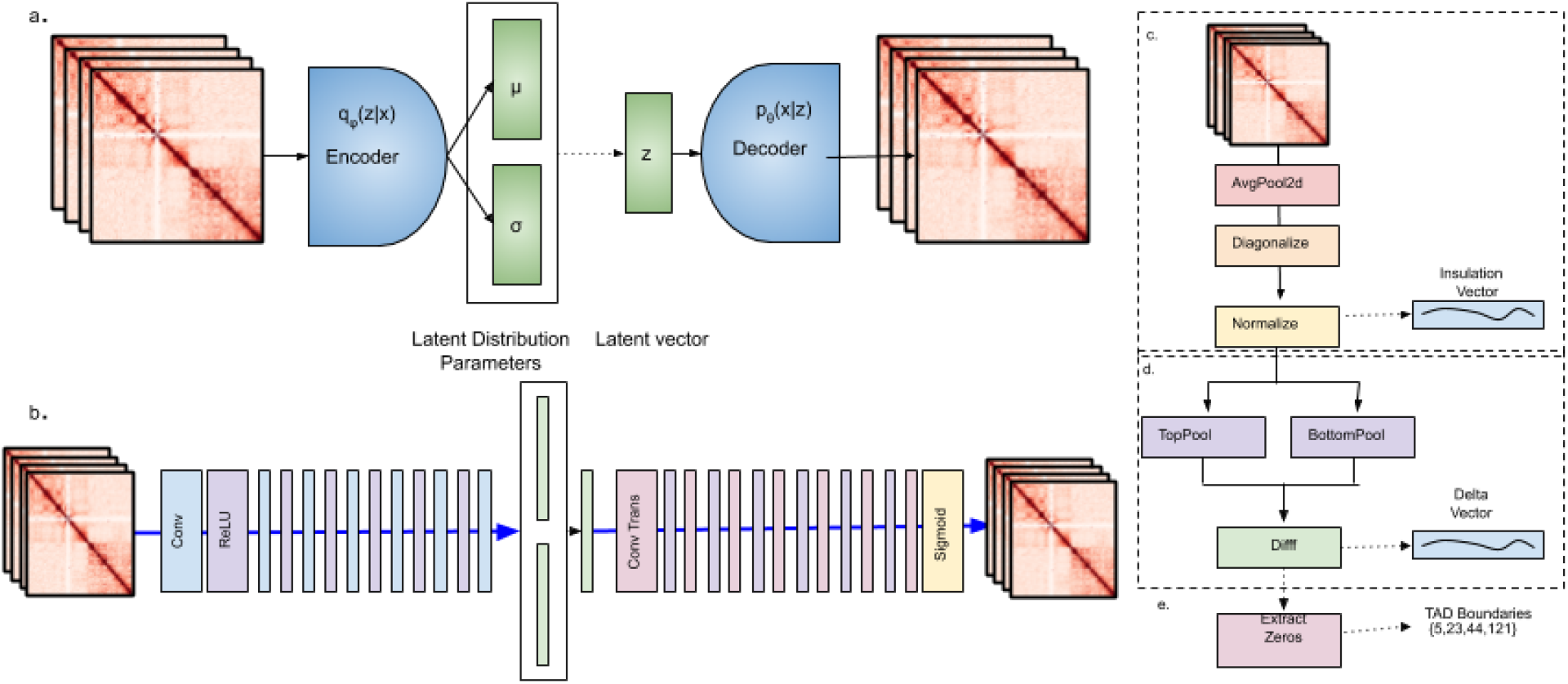
Variational Autoencoder (a) Overview of Variational Autoencoder Approach (b) VEHiCLE architecture. Tad loss evaluated using a feedforward implementation of Insulation loss computing (c) Insulation Vector (d) Delta Vector and (e) Identification of TAD Boundaries.

We compute variational loss by passing both the enhanced Hi-C contact matrix and target high resolution Hi-C contact matrices through the packpropagatable encoder component of our variational autoencoder network, extracting latent dimensional representations. We then compute the mean differences between their latent feature vectors,

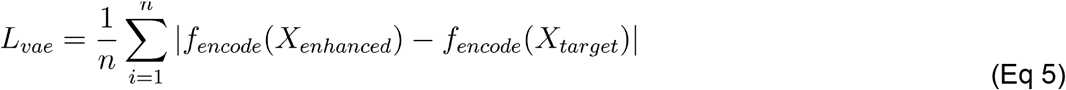

Where *f_encode_* is the encoding function, defined by our trained encoder network.

#### Insulation Score Loss

Most of the previously proposed loss functions for developing Hi-C enhancement networks draw upon loss functions prolific in the fields of computer vision ^5–9^. While there are certainly advantages to these strategies, they derive from assumed similarities between the tasks of image superresolution and Hi-C superresolution. However the tasks are not synonymous. Hi-C contact matrices contain important information used for downstream feature analysis such as loop calling, TAD identification and 3D model construction. Consequently images which are highly visually similar but which are blurry, shift positions of structural features, or contain noise might result in significant differences in downstream analysis. With this fact in consideration we used domain knowledge of computational genomics to devise an insulation loss function, which directly trains networks to correctly identify downstream features, specifically TAD placement.

One well established strategy for the identification of TADs is the use of insulation scores^12^. Insulation scores of a matrix are calculated by sliding a 50bin (500kb ×500kb) window down the diagonal of a matrix and summing the signal across each bin, resulting in an insulation vector (fig 1c). This insulation vector is normalized by taking the log2 ratio of each bin’s insulation score and the mean of all insulation scores on the chromosome. From the insulation vector a delta vector is computed by observing the change in signal strength 100kb downstream and upstream of each bin on the insulation vector (fig2d). This delta vector is treated as a pseudo-derivative, and identifies insulation valleys in the regions where the delta vector crosses the x-axis from negative values to positive values, indicating a relative minimum in insulation. TAD Boundaries are assigned to each insulation valley whose difference in strength between the nearest left local max and right local min was >0.1 (fig2e).

The insulation TAD calling procedure can be encoded into a single, back propagatable network up until extraction of the delta vector (fig2cd). We define insulation loss,

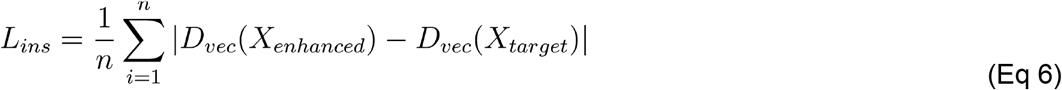

Where *D_vec_* is a backpropagable network which maps a contact matrix to a delta insulation vector.

#### Bin-Wise Mean Squared Error Loss

Bin-wise mean square error loss is a thoroughly tested loss function used in previous Hi-C enhancement literature ^8 6,9^. It contributes to maintaining visual similarity between enhanced and target Hi-C contact matrices.

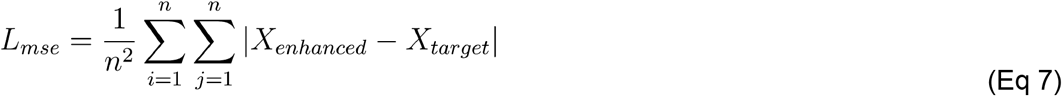

#### Composite Training Function

To capitalize on the advantages of all four loss functions we incorporate them into our comprehensive training process. First the variational network is trained on the train and validation datasets. Then the trained encoder is used for *L_vae_* along with the three other training losses to train the generator network, yielding our overall loss function

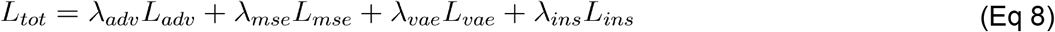

Where λ_*x*_ are hyperparameters used to determine loss contribution. We λ_*adv*_ = 0.0025, λ_*mse*_ = 1, λ_*vae*_ = .01, and λ_*ins*_ = 1.

## Results

### Latent space representations permit generation of synthetic Hi-C Data

The KL divergence term in the loss function of our variational autoencoder imposes constraints on the latent dimension, pushing our estimate for the prior *q*(*z*|*x*) to be close to a vector distribution of gaussian random variables. Because all latent vector variables fall within gaussians centered around 0, most vectors near the center of these gaussians can be successfully decoded into Hi-C space, resulting in a generative model for Hi-C data. We first perform PCA on our training set’s learned, latent dimensional features. We keep the first 15 principal components and create a function mapping PC values to the latent dimensional space. We then use our trained decoder network to transform the values in latent dimensional space into Hi-C space (figure 3a). The result is a function mapping 15 variables to 2.5Mb blocks of Hi-C data. We hook this function into an interactive matplotlib widget, permitting manual visualization of changes to generated Hi-C data as input variables are adjusted. The zero vector results in a vanilla Hi-C map with interaction frequency between two regions following the inverse of genomic distance (figure 3b). We observe that many of the tunable feature vector components correspond directly with biologically meaningful features in Hi-C space such as: formation of TADS, increasing TAD size (figure 3c), increasing TAD frequency, shifting TAD position (figure 3cd), formation of genomic stripes (figure 3e) ^13^ and formation of chromatin loops ^14^ (figure 3f).

**Figure 3.**
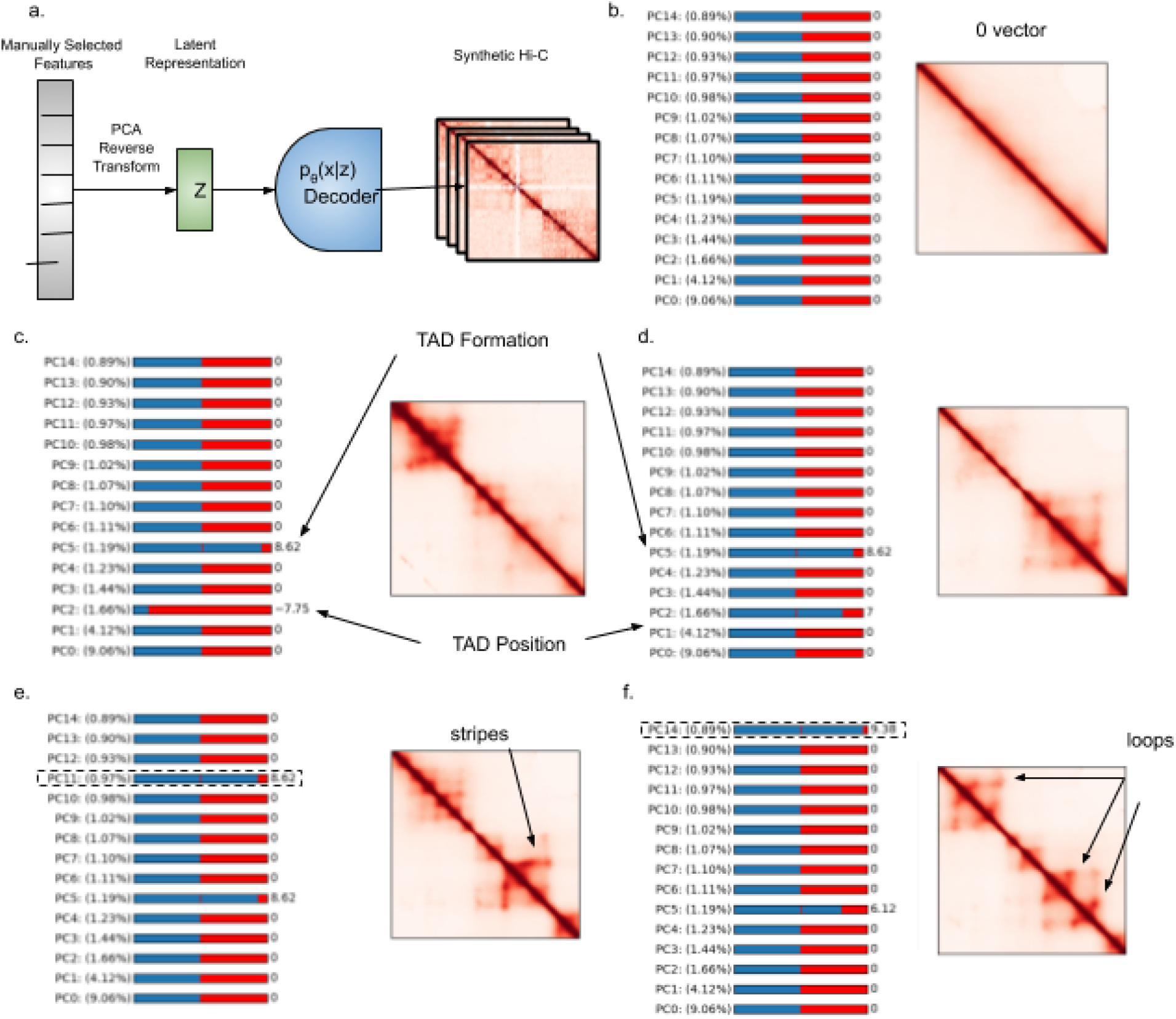
(a) Diagram of synthetic Hi-C generation tool, a user tunable zero-centered feature vector is transformed via PCA reverse transform to latent space and then passed through our tuned decoder network. (b) The 0 vector corresponds to a purely linear contact map. (c) Increasing value of PC5 results in generation of TADS (d) adjusting value of PC2 shifts position of TADs (e) Adjusting PC11 creates stripes within TADS (f) adjusting PC14 develops loops within TADs.

### Low Resolution Hi-c contact matrices enhanced by VEHiCLE appear visually competitive with other Enhancement algorithms

We generate visual heatmaps of Hi-C contact map reconstructions using VEHiCLE as well as three other previously developed algorithms: HiCSR, DeepHiC and HiCPlus. We observe high visual similarity between reconstructions by VEHiCLE and other enhancement algorithms (figure 4a.) We also subtracted High resolution contact maps from reconstructions by each tool to observe a visual difference matrix (figure 4b). Visually VEHiCLE appears competitive with existing algorithms.

**Figure 4.**
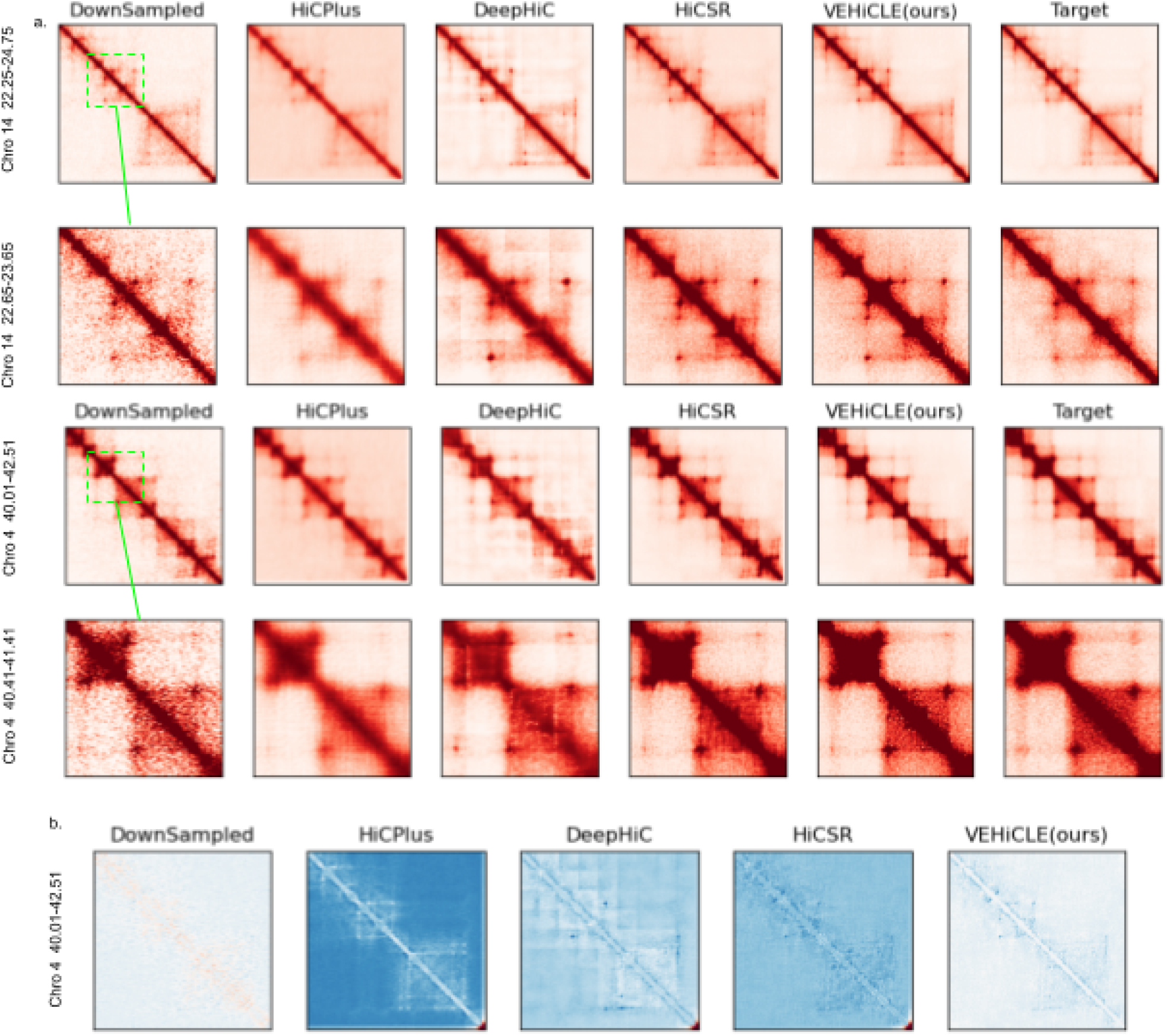
Visual Comparison of deep learning based methods for contact matrix enhancement. (a) Visual comparison of enhancement matrices. (b) Absolute difference matrices between target high resolution data and enhancement.

### Low Resolution Hi-C contact matrices enhanced by VEHiCLE achieve strong similarity to high resolution contact matrices using vision metrics

We evaluated the effectiveness of VEHiCLE in predicting high resolution contacts using 5 common metrics: Pearson Correlation Coefficient (PCC), Spearman Correlation Coefficient(SPC), Mean Squared Error (MSE), Signal-to-noise ratio (SNR) and Structure Similarity Index (SSI) (see methods.) We compared VEHiCLE reconstructions to the lower resolution data as well as other super resolution methods (HiCPlus, DeepHic and HiCSR.) VEHiCLE enhanced contact matrices consistently showed improvement relative to low resolution data along all 5 metrics (table 1). VEHiCLE frequently out-performed other Hi-C superresolution methods, obtaining the highest PCC and SPC scores for every test chromosome (table 1). VEHiCLE performed better than all models in every metric for chromosome 4 and performed better than DeepHiC and HiCPlus in every visual metric on all test chromosomes (table 1).

**Table 1.** Comparison of visual metrics across different super-resolution algorithms

| **Chro 4** | Downsampled | HiCPlus | DeepHiC | HiCSR | VEHiCLE |
| --- | --- | --- | --- | --- | --- |
| PCC | 0.7592 | 0.9103 | 0.9212 | 0.9285 | **0.9427** |
| SPC | 0.6259 | 0.805 | 0.7715 | 0.8292 | **0.8734** |
| SSIM | 0.2336 | 0.3284 | 0.3784 | 0.4346 | **0.4605** |
| MSE | 0.0468 | 0.0163 | 0.0162 | 0.0114 | **0.0092** |
| SNR | 306.65 | 514.24 | 516.847 | 619.41 | **693.967** |
| **Chro 14** | Downsampled | HiCPlus | DeepHiC | HiCSR | VEHiCLE |
| PCC | 0.8682 | 0.9374 | 0.9481 | 0.9583 | **0.9632** |
| SPC | 0.6692 | 0.8159 | 0.7188 | 0.85 | **0.89** |
| SSIM | 0.3524 | 0.4022 | 0.5481 | 0.5877 | **0.5987** |
| MSE | 0.0145 | 0.0117 | 0.0052 | **0.0041** | 0.0051 |
| SNR | 341.5712 | 380.0418 | 554.9034 | **627.7526** | 570.9596 |
| **Chro 16** | Downsampled | HiCPlus | DeepHiC | HiCSR | VEHiCLE |
| PCC | 0.8798 | 0.9327 | 0.9479 | 0.9602 | **0.9615** |
| SPC | 0.6684 | 0.8097 | 0.6949 | 0.8496 | **0.8875** |
| SSIM | 0.3901 | 0.3935 | 0.5618 | 0.5924 | **0.596** |
| MSE | 0.0118 | 0.0124 | 0.0047 | **0.0036** | 0.0053 |
| SNR | 332.8447 | 318.1057 | 517.1062 | **597.73** | 496.1329 |
| **Chro 20** | Downsampled | HiCPlus | DeepHiC | HiCSR | VEHiCLE |
| PCC | 0.91 | 0.93 | 0.95 | **0.97** | **0.97** |
| SPC | 0.69 | 0.83 | 0.69 | 0.86 | **0.9** |
| SSIM | 0.44 | 0.41 | 0.6 | **0.64** | 0.6 |
| MSE | 0.0076 | 0.0124 | 0.0038 | **0.0027** | 0.0054 |
| SNR | 364.83 | 282.43 | 510.35 | **608.04** | 432.403 |

**Table 2.** Comparison of Hi-C Superresolution algorithms on Hi-C specific Metrics

| **Chro 4** | GenomeDISCO | HiCRep | QuASAR-Rep |
| --- | --- | --- | --- |
| Down Sampled | 0.941 | 0.967 | 0.924 |
| HiCPlus | 0.972 | 0.974 | 0.995 |
| DeepHiC | 0.945 | 0.972 | 0.993 |
| HiCSR | **0.98** | 0.989 | **0.995** |
| VEHiCLE | 0.979 | **0.99** | **0.995** |
| **Chro 14** | GenomeDISCO | HiCRep | QuASAR-Rep |
| Down Sampled | 0.942 | 0.982 | 0.944 |
| HiCPlus | 0.933 | 0.969 | 0.995 |
| DeepHiC | 0.907 | 0.97 | 0.993 |
| HiCSR | **0.979** | **0.991** | **0.996** |
| VEHiCLE | 0.965 | 0.986 | **0.996** |
| **Chro 16** | GenomeDISCO | HiCRep | QuASAR-Rep |
| Down Sampled | 0.927 | 0.974 | 0.941 |
| HiCPlus | 0.904 | 0.948 | 0.992 |
| DeepHiC | 0.88 | 0.96 | 0.99 |
| HiCSR | **0.972** | **0.987** | **0.994** |
| VEHiCLE | 0.955 | 0.976 | **0.994** |
| **Chro 20** | GenomeDISCO | HiCRep | QuASAR-Rep |
| Down Sampled | 0.934 | 0.981 | 0.955 |
| HiCPlus | 0.895 | 0.949 | 0.994 |
| DeepHiC | 0.864 | 0.959 | 0.99 |
| HiCSR | **0.974** | **0.988** | **0.996** |
| VEHiCLE | 0.948 | 0.979 | **0.996** |

### Downsampled Hi-C contact matrices enhanced by VEHiCLE display Significant improvement using Hi-C specific metrics

We next evaluated VEHiCLE reconstructions using 3 Hi-C specific metrics: GenomeDISCO, HiCRep and QuASAR-Rep (see methods). We observe consistent improvement in VEHiCLE enhanced matrices relative to down sampled matrices across all 3 metrics. VEHiCLE enhanced metrics remain competitive with other methods, beating DeepHiC and HiCPlus in all instances and tying with HiCSR on QuASAR-Rep (table 3). These results indicate biological consistency with VEHiCLE enhanced matrices.

**Table 3.** L2 norm of TAD Insulation difference vectors

|  |  |  |  |  |  |
| --- | --- | --- | --- | --- | --- |
| Norm of<br>Insulation Score<br>Difference<br>Vectors |  |  |  |  |  |
| **Chr 4** | Downsampled | HiCPlus | DeepHiC | HiCSR | VEHiCLE |
| GM18278 | 7.966 | 6.64 | 7.52 | 4.389 | **3.996** |
| K562 | 10.942 | 9.1133 | 9.957 | 7.605 | **6.5769** |
| IMR90 | 9.8344 | 8.5681 | 9.2244 | 5.9457 | **4.5372** |
| HMEC | 16.143 | 13.132 | 15.267 | **10.17** | 10.685 |
| **Chr 14** | Downsampled | HiCPlus | DeepHiC | HiCSR | VEHiCLE |
| GM18278 | 2.68 | 2.619 | 3.774 | 2.898 | **2.456** |
| K562 | 6.225 | 5.927 | 6.329 | 5.548 | **5.318** |
| IMR90 | 4.3609 | 4.6005 | 5.1827 | 3.871 | **3.439** |
| HMEC | 9.214 | 8.34 | 9.549 | **7.448** | 8.46 |
| **Chr 16** | Downsampled | HiCPlus | DeepHiC | HiCSR | VEHiCLE |
| GM18278 | 4.162 | 3.467 | 3.769 | 3.099 | **3.01** |
| K562 | 6.653 | 5.903 | 6.485 | 5.14 | **4.976** |
| IMR90 | 5.806 | 5.0148 | 5.459 | 4.169 | **4.03807** |
| HMEC | 8.957 | 8.353 | 8.799 | **7.527** | 7.806 |
| **Chr 20** | Downsampled | HiCPlus | DeepHiC | HiCSR | VEHiCLE |
| GM18278 | **2.077** | 2.419 | 2.587 | 2.624 | 2.337 |
| K562 | 5.316 | 4.835 | 5.021 | **4.307** | 4.396 |
| IMR90 | 2.888 | 3.522 | 3.444 | 3.083 | **2.746** |
| HMEC | 6.383 | 6.662 | 6.579 | **5.805** | 6.202 |

### VEHiCLE Enhanced contact matrices effectively retrieve downstream features such as TADS

We identified TADs using the prolific insulation score method ^12^. This method assigns an insulation score vector by sliding a window across the diagonal of the contact matrix, constructing an insulation difference vector, and using the zeros of the insulation difference vector to discover TAD boundaries (see methods). We compare the insulation difference vector of each matrix-enhancement algorithm to the insulation difference vector of our high resolution contact matrix using the L2 norm dissimilarity metric. In most cases VEHiCLE enhanced insulation difference vectors have higher similarity to target matrices relative to other matrix enhancing algorithms. (Table 3). Furthermore, even in instances where VEHiCLE is outperformed by another algorithm we consistently observe higher similarity between the target high resolution matrices and VEHiCLE enhanced matrices relative to low resolution matrices. (figure 5a, Table 3).

**Figure 5.**
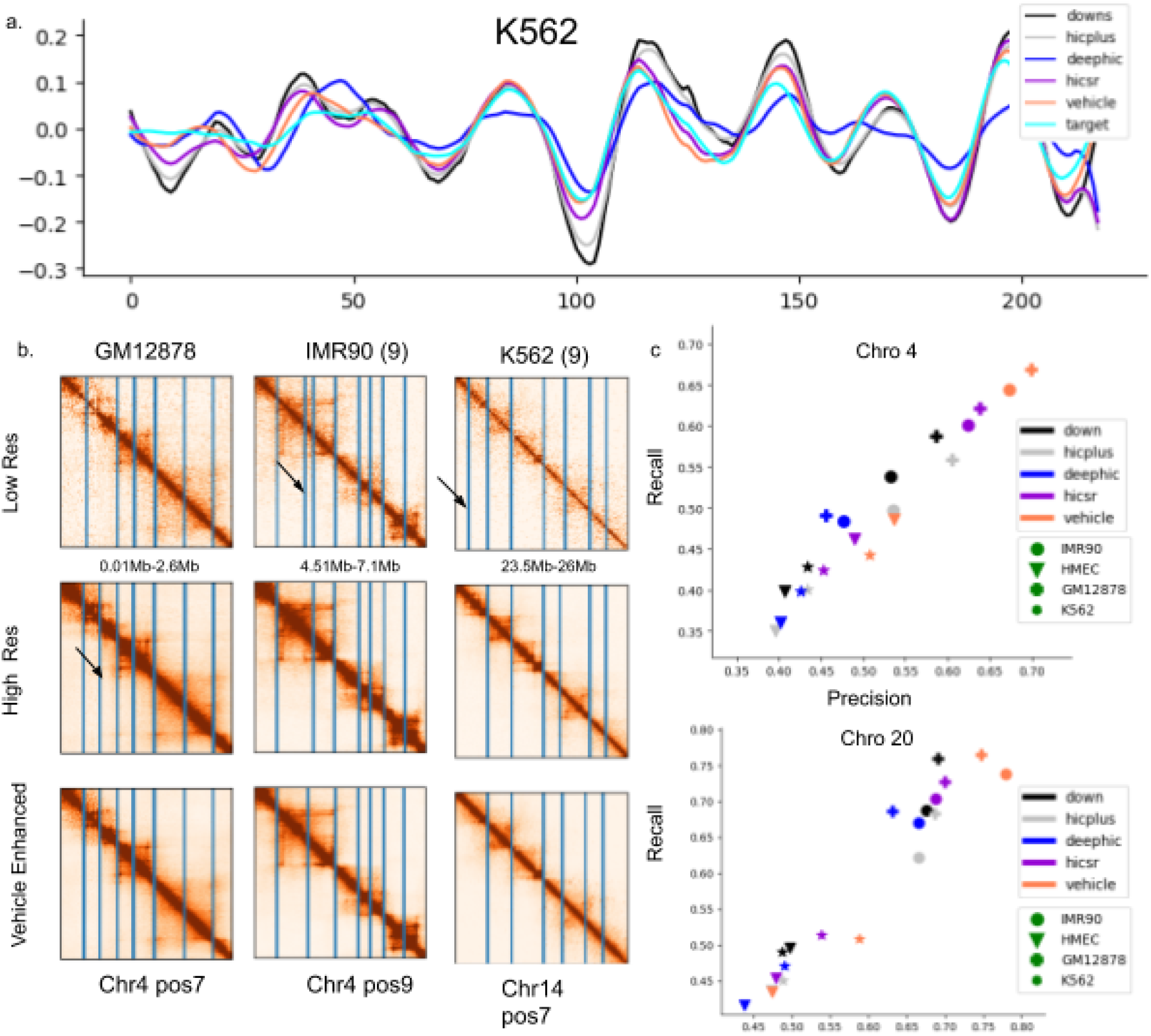
TAD Analysis (a) Insulation difference vector for Downsampled, Target and algorithmically enhanced matrices. (b) Missing TAD’s are recovered and Faulty TADs are removed in VEHiCLE enhanced matrices. (c,d) Precision Recall Values of Algorithmically enhanced TAD boundaries across cell lines.

### 3D Chromatin Model Construction

We tested the effectiveness of reconstructed data in building 3DModels using the structural modeling tool 3DMax. We extracted constraints from the low resolution, high resolution and VEHiCLE-enhanced 2.57Mb×2.57Mb regions of our test dataset chromosomes of the GM12878 dataset. From each constraint grouping we generated 3 models. We observed significantly higher visual similarity between VEHiCLE-enhanced and high-resolution matrices relative to low-resolution matrices (Fig 6a). We then used the TM-score metric to quantify structural similarity of models. We observed higher TM-scores between high resolution and VEHiCLE-enhanced matrices than between high resolution and low resolution models (Fig 6b). We also observed higher TM-score similarities between models generated by the same VEHiCLE-Enhanced matrices relative to models generated by the same low resolution matrices, indicating VEHiCLE enhanced models are more consistent (Fig 6c).

**Figure 6.**
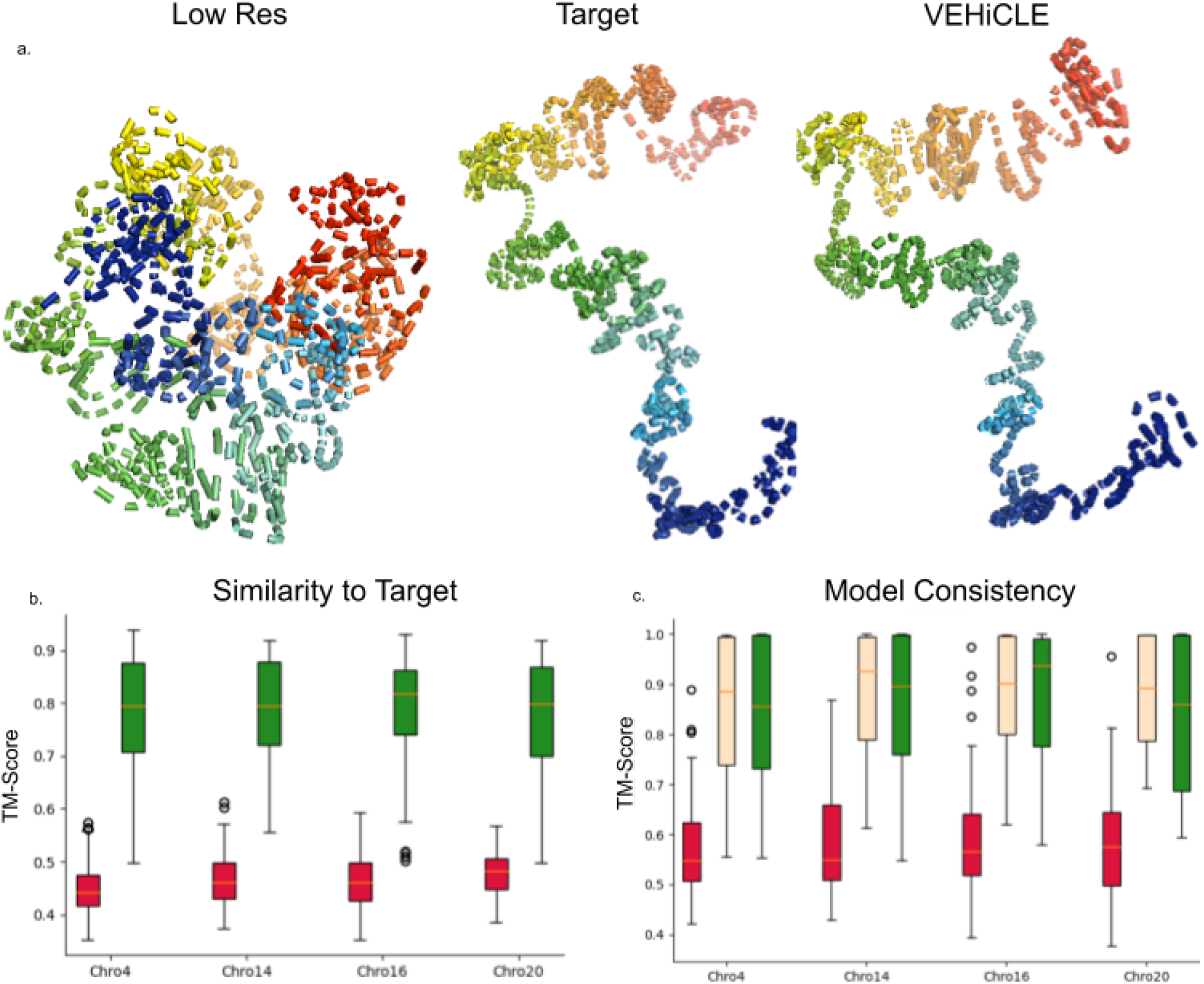
(a) 3D reconstruction of Chro 20 0.6MB-3.1MB. (b) TM-score comparison of High Resolution structures to (red) Low resolution structures and (green) VEHiCLE enhanced structures. VEHiCLE enhanced scores are significantly higher (wilcoxon rank sum p value < 1e-20) (c) Average TM-Score comparison of ingroup structures generated by same contact matrix (red) low res, (yellow) high res, (green) VEHiCLE enhanced. VEHiCLE enhanced scores are significantly better than low-resolution scores (wilcoxon rank sum p value < 1e20)

## Discussion

One of the most common challenges in Deep Learning projects is the opaque nature of a neural network’s inner functioning. Consequently our ability to extract latent features and map them to biologically relevant structures provides a significant advance in increasing interpretability of Hi-C matrices. Our GUI tool can be used to generate Hi-C data through user tunable parameters with biologically relevant downstream structures such as TAD strength, TAD positioning, stripes and loops. Further inspection of these features have potential to further analysis of key characteristics of chromatin organization

Our introduction of the Insulation loss sets a new precedent of utilizing biological knowledge in the training of Hi-C networks. This may open the door for future improvement of Hi-C data enhancement by utilizing other forms of domain knowledge to increase usability of deep learning enhanced matrices. Future loss functions could incorporate algorithms for identification of other important downstream features such as loops or stripes.

In addition to the increased interpretability and inclusion of domain knowledge, VEHiCLE obtains resolution enhancement results competitive with the state-of-the art, often beating top algorithms on a variety of metrics, all while preserving the ability to convey meaningful structures such as TAD’s and 3D structure in downstream analysis.

## Methods

### Dataset Assembly

Like many of the previous Hi-C super resolution networks we train VEHiCLE on high and low resolution Hi-C data for the GM12878 cell line ^15^. While previous work often split chromosomes into training, validation and testing sets in a sequential manner ^8 9^ we were concerned that differences in the 3D conformation of large vs small chromosomes ^16^ may contain implicit bias in contact map features that could confound training. Consequently we assembled training, validation and test sets in a non sequential manner using chromosomes 1,3,5,6,7,9,11,12,13,15,17,18,19,21 as our training set, chromosome 2,8,10,22 as our validation set and chromosomes 4,14,16,20 as our test set.

Previous work on Hi-C super resolution consistently used network input window sizes of 0.4Mb × 0.4Mb at 10kb resolution, requiring networks to split chromosome contact maps into 40×40bin matrices ^5–9^. While this strategy has seen relative success, a major disadvantage is that certain important features of Hi-C such as TADs can span ranges larger than 0.4Mb, meaning that it is impossible for previous networks to explicitly encode important information about TAD organization. Furthermore this informational bottleneck of constraining window sizes to 40×40 bins is not incumbent upon the employed super-resolution networks as work in the field of computer vision has demonstrated the effectiveness of GAN and VAE networks on significantly larger images. With these considerations in mind we instead built our network to accept 2.69 Mb × 2.69Mb images, a range which is large enough to fully encompass the average TAD of length 1MB ^17^. Observing 2.69Mb × 2.69Mb regions of Hi-C contact maps at range 10kb results in submatrix images of 269×269 bin size. Because of the expanded window size we trained our network exclusively on diagonally centered submatrices, split by sliding a 269×269 window down the diagonal of each chromosomes Hi-C contact map resulting in a total of 3309 training, 1051 validation, and 798 testing matrices.

Models were only trained using the GM12878 Cell line. Experiments were run on the test chromosomes of K562, IMR90 and HMEC cell lines to verify the effectiveness of our network at retrieving information when trained on a different cell line.

#### Variational Autoencoder architecture

The VAE component of VEHiCLE utilizes two neural networks for the encoding and decoding components, where the encoder is trained for the parameters of *q_θ_* and the decoder is trained to optimize the parameters of *p_θ_*. The VEHiCLE encoder network contains 7 convolutional layers with kernel counts: 32,64,128,256,256,512,512. Each convolutional layer is separated by leaky ReLU and batch normalization. The Decoder network has 7 layers of Convolution Transpose with the kernel counts 512, 512, 256, 256, 128, 64, 32, also separated by leaky ReLU and batch norm functions. The Decoder network is appended by a Sigmoid activation function placing outputs in the range of [0,1].

#### Generative Adversarial Network Architecture

We use the Discriminator and Generator architecture defined in HiCSR, with the exception of our generators final non-linearity, which is changed from tanh, to a sigmoid so that outputs are mapped to [0,1]. The generator architecture contains 15 residual blocks separated by skip connections, each containing 64 convolutional filters. The fully convolutional Discriminator is a fully convolutional network with ReLU activation. Both the generator and discriminator are trained with Batch Normalization

#### Other Networks

We used the pytorch versions of HiCPlus, DeepHiC and HiCSR provided at https://github.com/wangjuan001/hicplus, https://github.com/omegahh/DeepHiC and https://github.com/PSI-Lab/HiCSR. We first tested networks using their literature provided weights, however we obtained very poor performance because these networks were trained on alternative training sets with key characteristic differences from ours. First, their training sets had bin value ranges of [−1,1], however our training datas range was [0,1] because negative values confound the probabilistically motivated VAE component. Second the input size of contact maps for previous networks was 40×40, while our network aims to incorporate surrounding genomic information and utilizes a larger window input size (269×269). To provide more accurate comparison we trained networks on our own GM12878 Dataset. Because our networks accept a large scale input matrix 269×269, but other networks were built to accept 40×40 pieces, we trained other networks by splitting each 269×269 into 36 non-overlapping pieces. Evaluation of Hi-C metrics was performed by feeding split pieces through networks as necessary, then reassembling pieces and comparing full chromosome contact maps.

### Visual Metrics

We utilize 5 reproducibility metrics pulled from image-super resolution literature: Pearson Correlation Coefficient (PCC), Spearman Correlation Coefficient(SPC), Mean Squared Error (MSE), Signal-to-noise ratio (SNR) and Structure Similarity Index (SSI)

#### Mean Squared Error

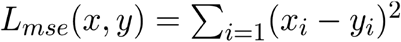

#### Pearson Correlation Coefficient

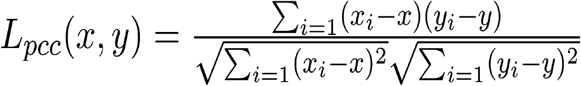

#### Spearman Correlation Coefficient

Spearman Correlation is similar to pearson correlation differing in that it utilizes rank variables so as to evaluate monotonic relationship between the matrices without imposing a linearity condition that may not exist in nature.

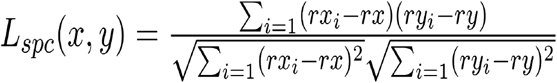

#### Signal-To-Noise Ratio

Signal-To-Noise Ratio uses a ratio of the clean signal to the difference between clean and noisy signals to represent how much signal is actually getting through. The higher the value of SNR the better quality the data.

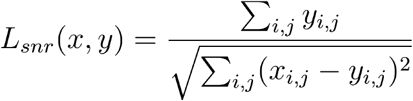

#### Structural Similarity Index

SSI is calculated by sliding windows between images and averaging values. The constants *C_1_* and *C_2_* are used to stabilize the metric while the means, variances and covariances are computed via a Gaussian filter. We use the implementation of SSI developed by Hong et al^8^ (DeepHiC) keeping their default values for the size of sub-windows and variance value of gaussian filter at 11 and 3 respectively.

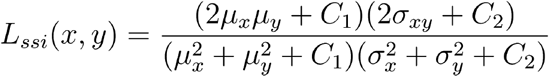

### Hi-C Reproducibility Metrics

We consider 3 Hi-C specific reproducibility metrics: GenomeDISCO^18^, HiCRep ^19^, and QuASAR-Rep^20^. We use the 3DChromatin_ReplicateQC ^21^ implementations of the metrics. This 3DChromatin_ReplicateQC repository also included metrics for the tool HiC-Spector ^20^, however we consistently obtained faulty values, even when using the repositories sample data and so we excluded HiC-Spector results from this analysis. GenomeDISCO utilizes a random walk on a graph generated by contact maps to obtain a concordance score ^21^. HiCRep develops a stratum adjusted correlation coefficient for matrix comparison by measuring weighted similarity of contacts in identified stratum^21^. QuASAR-Rep calculates a correlation matrix of interaction using weights based on enrichment ^20^.

### Topologically Associated Domain Identification

Topologically associated domains were identified using Insulation score as identified in Crane et al. We mimicked their procedure entirely with the exception that our initial insulation score window size was condensed to 20 bins instead of 50 because this demonstrated greater visual accuracy in TAD positioning. ^12^

### Three Dimensional Model Reconstruction

To generate models we utilize 3DMax ^22^ with out of the box parameters of 0.6 conversion factor, 1 learning rate and 10000 max iteration. We create 3 models per input contact matrices. We generate models for every 5th 269Mb×269Mb input matrix from our training dataset, because this skipping distance ensures coverage of each chromosome while minimizing model generation time. Similarity between structures was measured using TM-score ^23^

## Data availability

All Hi-C data were downloaded from the Gene Expression Omnibus (GEO) GSE63525. For the Hi Resolution Matrices of GM12878, IMR90, K562 and HMEC we used GSE63525_GM12878_insitu_primary+replicate_combined_30.hic, GSE63525_IMR90_combined_30.hic, GSE63525_K562_combined_30.hic and GSE63525_HMEC_combined_30.hic respectively. For low resolution matrices we used GSM1551550_HIC001_30.hic, GSM1551602_HIC053_30.hic, GSE63525_K562_combined_30.hic, and GSM1551610_HIC061_30.hic respectively.

## Code Availability

VEHiCLE was built using python. All experimental code as well as the VEHiCLE enhancement tool and Contact Matrix generating GUI are available at https://github.com/Max-Highsmith/lsdcm.

## Acknowledgements

This project is partially supported by two NSF grants (no. IOS1545780 and no. DBI1149224).

## Author Contributions

MH and JC conceived the project. MH performed all experiments and drafted the manuscript. JC revised the manuscript.

## Competing Interests

The authors declare no competing interests

## Notes

### Competing Interest Statement

The authors have declared no competing interest.

